# Sex determination of Syrian hamster pups using placental tissues

**DOI:** 10.1101/2025.11.22.689895

**Authors:** Yana Kumpanenko, Lindsey Piessens, Victor Neven, Kai Dallmeier, Yeranddy A. Alpizar

## Abstract

Molecular sex determination in Syrian hamsters (*Mesocricetus auratus*) has been limited by incomplete annotation of Y-linked loci in currently available genome assemblies. Here, we evaluate the Y-linked gene *PRSSLY*, which encodes a testis-specific serine protease-like protein, as a molecular marker for genetic sexing of Syrian hamster embryonic and placental tissues. Primers flanking a conserved *PRSSLY* coding region produced a male-specific amplicon showing 100% concordance with results from the established *KDM5C/KDM5D* PCR assay in E15.5 tail biopsies. SYBR Green–based qPCR enables the accurate detection of *PRSSLY*, characterized by a unique melt-curve profile, exclusively in male samples, allowing for efficient and sensitive mid-throughput analysis. Among 417 placental samples from 39 dams tested, which naturally contain a mixture of maternal and embryonic genomes, the sex of individual pups was classified with 85% specificity, with the remaining ambiguous samples confirmed as male by PCR and agarose gel electrophoresis. This assay provides a robust and reproducible approach for accurate sex genotyping in developmental and reproductive studies using Syrian hamsters.

## INTRODUCTION

Genotyping methods for sex identification typically rely on the detection of Y-linked genes, whose presence distinguishes males from females. Several Y-linked genes are highly conserved among placental mammals, including *ZFY, SRY, TDY, KDM5*, and *PRSSLY* [1, 2]. In particular, the zinc-finger genes (*ZFX* and *ZFY*) and the sex-determining region Y gene (*SRY*) have been widely used for molecular sex identification across diverse taxa. A single primer set targeting conserved regions in these genes enables accurate sex determination in multiple mammalian species, including various mice and voles, bats, pigs, horses, cats, dogs, beluga whales, and humans, among others [3-6]. For Syrian hamsters (*Mesocricetus auratus*), which constitute an emerging small animal model for infectious [7, 8] and non-communicable diseases [9], sexing protocols have not been well established.

*KDM5* has previously been exploited to develop a qPCR-based assay for sexing Syrian hamster fetal cell cultures [10]. *KDM5* encodes lysine demethylase 5, a histone H3K4 demethylase that occurs as distinct isoforms on the X and Y chromosomes. The X-linked isoform (*KDM5C*) and its Y-linked homolog (*KDM5D*) share high sequence similarity but differ in gene structure and length [1, 11], making them suitable targets for molecular sex determination.

However, the use of embryonic or placental tissues poses additional challenges. Depending on gestational stage and tissue composition, male-derived cells may be underrepresented, leading to weak or inconsistent amplification of Y-linked targets. This limitation is particularly pronounced in placental tissue, where male fetal cells are mostly confined to the labyrinth layer, contributing only minimally to the overall genomic DNA yield and thereby diminishing assay sensitivity.

In this study, we evaluated *PRSSLY*, a Y-linked gene encoding a testis-specific serine protease-like protein that shows evolutionary conservation across mammalian species [2]. We demonstrate that *PRSSLY* can be reliably detected in placental tissue despite the predominance of maternal DNA, a context in which *KDM5D*-based assays fail to yield consistent results. These findings establish *PRSSLY* as a molecular target for accurate sex genotyping of Syrian hamster embryonic and placental tissues.

## METHODS

### Study approval

Animal experiments were approved by the KU Leuven ethical committee (project numbers P165/2022, P120/2025, and P088/2024), conducted following Federation of European Laboratory Animal Science Associations (FELASA) guidelines.

### Animals

Wild-type Syrian hamsters (*Mesocricetus auratus*, strain RjHan:AURA; 8-week-old males and females) were obtained from Janvier Laboratories and housed separately in individually ventilated cages (IsoCage N Biocontainment System, Tecniplast) under controlled conditions (21 °C, 55% humidity, 12:12 h light–dark cycle) with *ad libitum* access to food, water, and environmental enrichment (cardboard shelters and wooden blocks). Females were acclimatized for two weeks and mated overnight with fertile males during the proestrus phase, identified by copious vaginal secretions three days prior. Successful mating was confirmed the following morning (defined as embryonic day 0.5, E0.5) by the presence of sperm in vaginal smears. On embryonic day 15.5 (E15.5), females were euthanized by intraperitoneal injection of Dolethal (500 µL; 200 mg/mL sodium pentobarbital, Vétoquinol SA). Fetuses were collected by caesarean section. Tail and placental biopsies, latter ideally including sufficient tissue from the fetal side, were obtained for genotyping.

Wild-type C57BL/6 mice (*Mus musculus*; 8-10 weeks old) were obtained from Janvier Labs.

### Sequence alignment

Protein sequences of Y-linked serine-like proteases from *Peromyscus eremicus* (cactus mouse), *Sus scrofa* (domestic pig), *Tupaia chinensis* (Chinese tree shrew), *Mesocricetus auratus* (Syrian golden hamster), *Mus musculus* (house mouse), and *Microcebus murinus* (gray mouse lemur) were retrieved from NCBI (accession numbers DAZ89691.1, QBK17227.1, DAZ89689.1, URN45584.1, AIB55773.1, and DAZ89688.1, respectively).

Multiple sequence alignment was performed using ClustalW (https://www.genome.jp/tools-bin/clustalw) with default parameters. Gap penalties and substitution matrices were applied as defined by ClustalW defaults to ensure optimal alignment quality.

### Genomic DNA isolation and sex genotyping

Genomic DNA (gDNA) was isolated from hamster placental tissue (∼5-10 mg) and tail or ear biopsies (1–2 mm) using the Platinum™ Direct PCR Universal Master Mix (Thermo Fisher Scientific, Cat. No. A44647100) according to the manufacturer’s lysis protocol. Samples were incubated in Lysis Solution containing Proteinase K for 2 h at room temperature and heated to 98 °C for 1 min. Lysates were centrifuged, and DNA concentration was measured by spectrophotometry (NanoDrop). Two microlitres of lysate (150-200 ng DNA) were used directly as PCR template.

Quantitative PCR (qPCR) genotyping was performed using the iTaq™ Universal SYBR® Green One-Step Kit (Bio-Rad, Cat. No. 1725150) on a LightCycler® 96 platform (Roche Diagnostics). Reactions (20 µL) contained 10 µL master mix, 0.2 µL each of 1:10-diluted forward and reverse primers (**Table 1**), 0.25 µL reverse transcriptase, 7.35 µL nuclease-free water, and 2 µL gDNA. Cycling conditions were: 94 °C for 2 min; 40 cycles of 94 °C for 15 s, 55 °C for 15 s, and 68 °C for 20 s. Melt-curve analysis was performed immediately after qPCR with sequential 10 s holds at 0.5 °C increments from 65 °C to 95 °C.

**Table 1.**
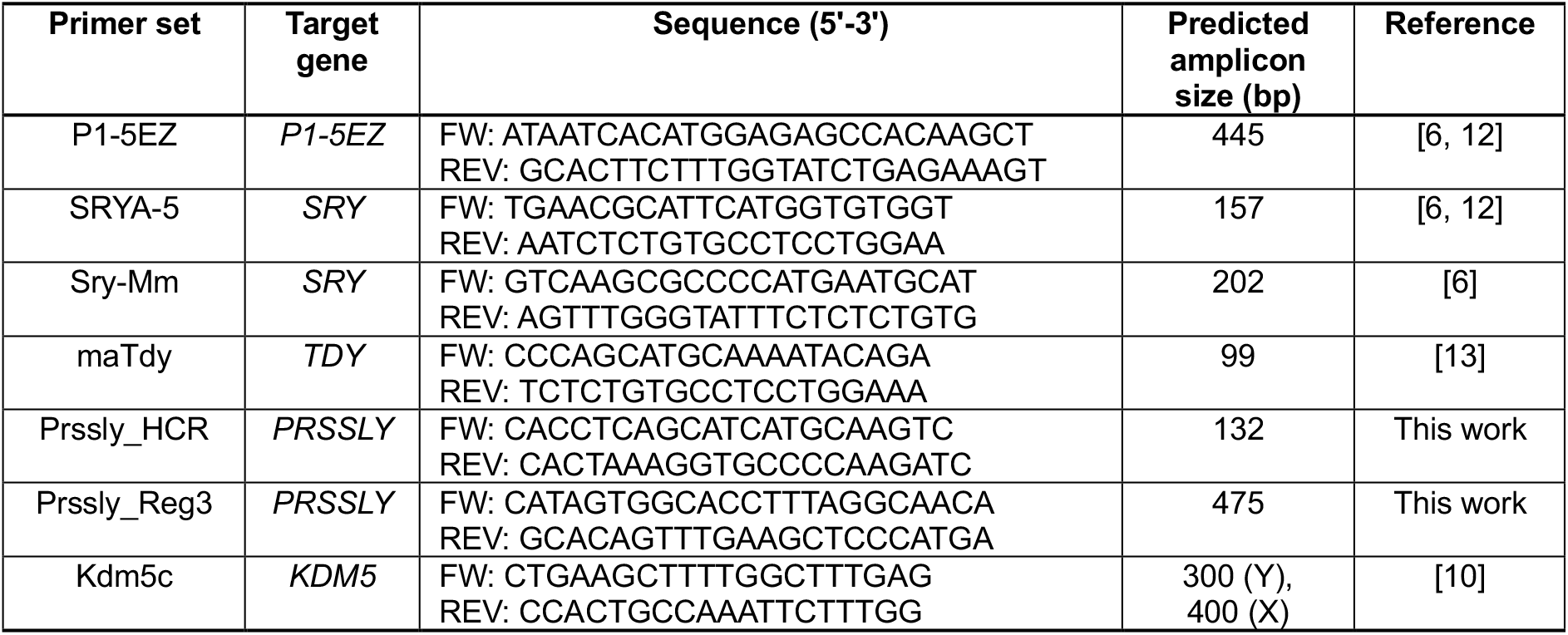
List of primers used in this study. All primers were synthesized and purchased from Integrated DNA Technologies (IDT).

Conventional PCR was performed using the Platinum™ Direct PCR Universal Master Mix (Invitrogen, Cat. No. A44647100) (10 µL master mix, 0.75 µL each of 1:10-diluted forward and reverse primers, 6.5 µL nuclease-free water, and 2 µL gDNA) and the cycling as described for the qPCR assay. PCR products were resolved by electrophoresis on 1.5% agarose gels in 1× TAE buffer containing Midori Green, with a 10 µL DNA ladder (Promega, Cat. No. G2101) as reference.

PCR bands were excised and purified using the Monarch® Spin DNA Gel Extraction Kit (New England Biolabs, NEB #T1120L), following the manufacturer’s instructions. Sequencing of the purified PCR products was performed on an ABI 3730xl DNA Analyzer by Macrogen (Seoul, South Korea).

## RESULTS AND DISCUSSION

### Identification and validation of *PRSSLY* as a marker for sex identification in Syrian hamsters

Previously described universal primer sets directed at *SRY* and *TDY*, which discriminate sex in several other mammalian species [6, 12, 13], did not yield sex-specific PCR products in Syrian hamsters (**Supplementary Figure 1**). To identify a suitable marker, we designed primers spanning a conserved *PRSSLY* coding region with a predicted amplicon size of 132 bp (**Figure 1A** and **Table 1**). However, the PCR produced multiple fragments, including a prominent >1.5-kb band specific to males (**Figure 1B**).

**Figure 1.**
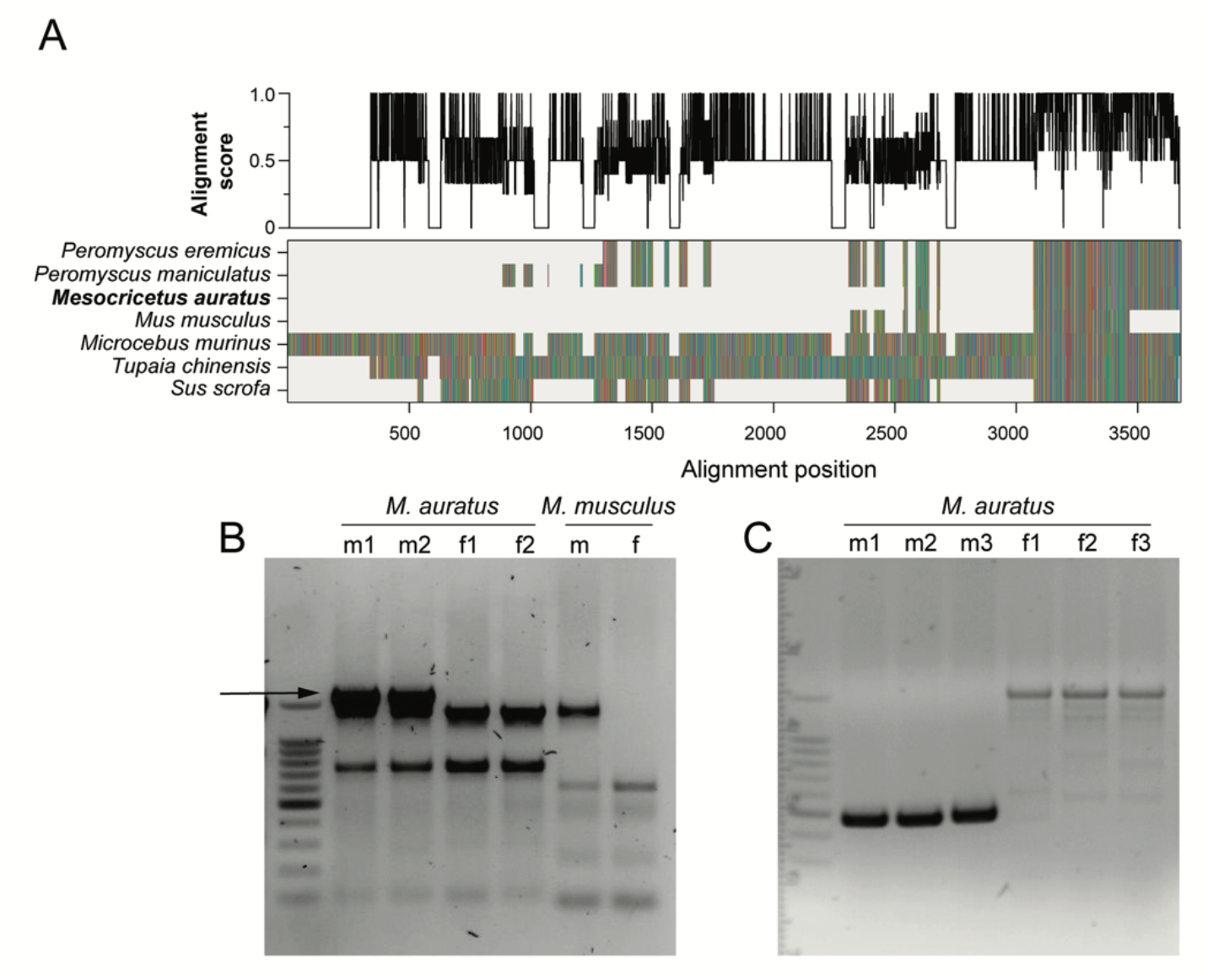
Identification of a conserved Y-linked serine protease region and primer validation. (**A**) Multiple sequence alignment of PRSSLY from representative mammalian species, with amino acids colour-coded according to physicochemical properties: hydrophobic (blue), aromatic (purple), polar uncharged (green), positively charged (red/pink), negatively charged (orange) and other (brown). Alignment gaps are shown in light grey. The upper panel displays the per-column conservation score from 0 (min) to 1 (max). A highly conserved region was selected for primer design. (**B**) PCR amplification using a primer set targeting the broadly conserved region (Prssly_HCR) on gDNA isolated from ear biopsies of adult hamsters and mice. A distinct male-specific band (arrow) was obtained, purified, and sequenced. (**C**) Refined amplification using the second-generation primer set (Prssly_Reg3), designed from the sequenced male-specific product in panel B. The optimized primers consistently produced a strong male-specific band of 475 bp, with only faint, nonspecific amplification in female samples.

Direct sequencing of this band revealed intronic sequences interrupting the predicted CDS locus. Guided by this genomic structure, we developed a revised primer pair, which eliminated most nonspecific amplification and yielded a single male-specific band of 475 bp; female samples generated only faint, nonspecific products of higher molecular weight (**Figure 1C**).

For mid-throughput screening, we next established a SYBR Green–based qPCR assay complemented by melt-curve analysis for increased specificity. In this format, robust amplification was consistently observed in male samples, with a mean Cq value of 25.6 ± 0.3 (n=13). The corresponding melting profiles were characterized by two distinct peaks at 76.5 °C and 79.0 °C. By contrast, female samples exhibited either no amplification or aberrant late amplification with a fluorescence detection delay of at least 13 cycles (mean Cq value 39.7 ± 0.8; n=13). Consequently, melting profiles in female samples were either indeterminable or exhibited inconsistent, non-reproducible patterns lacking the two characteristic peaks (**Figure 2A-C**).

**Figure 2.**
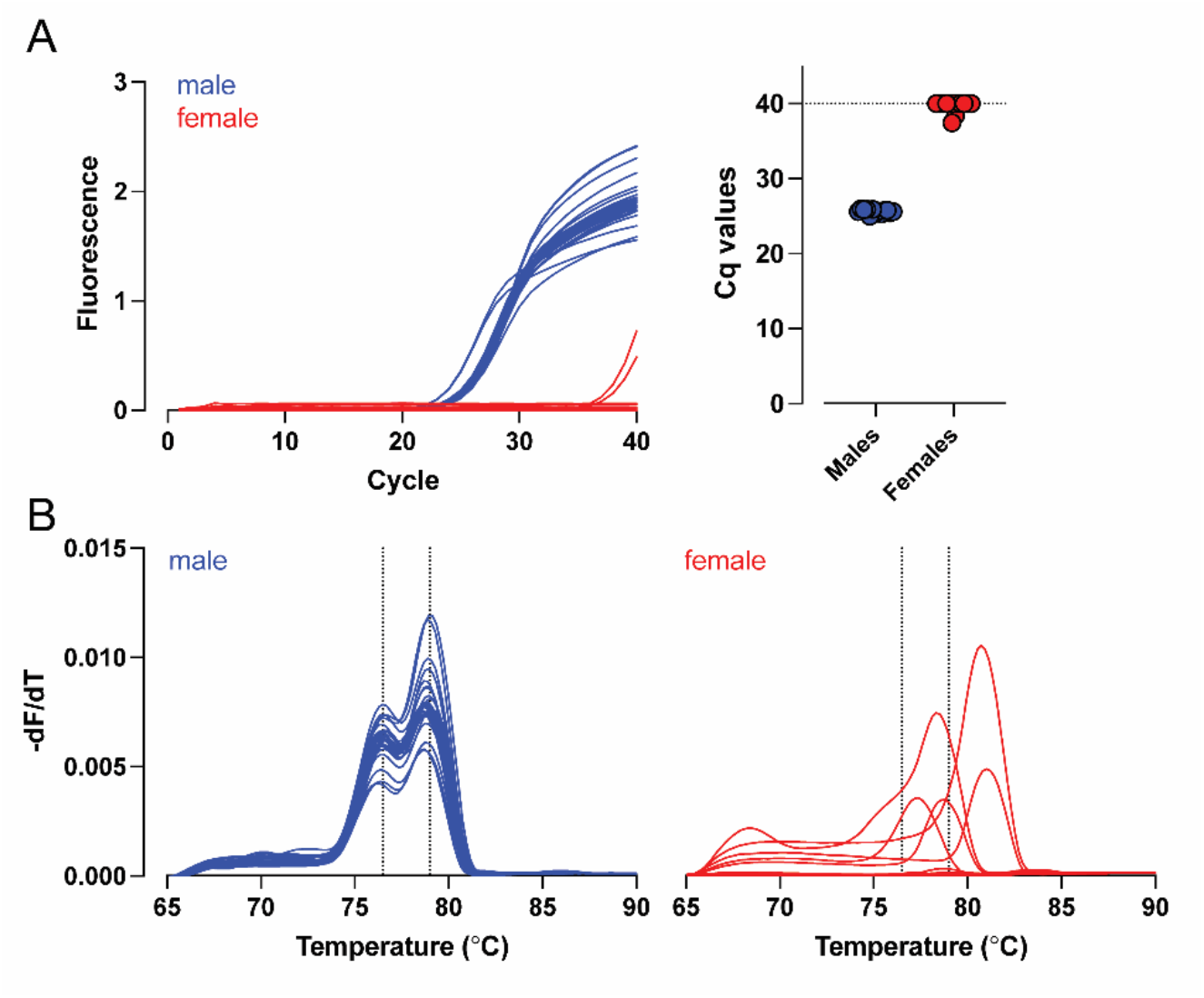
Validation of a SYBR Green–based qPCR assay for mid-throughput sex identification. (**A**) Real-time PCR amplification curves generated from gDNA isolated from ear biopsies of adult male (blue) and female (red) individuals. Male samples consistently exhibited lower Cq values, indicating specific amplification of the Y-linked target region, whereas female samples produced no amplification or only a delayed, nonspecific signal. (**B**) Melting curve analysis showing the temperature-dissociation profiles of the PCR products. Dotted vertical lines indicate the peak melting temperatures for male-specific amplicons at 76.5 °C and 79.0 °C.

### Sex identification in embryonic biopsies using *PRSSLY*-based qPCR

To assess the applicability of *PRSSLY* for embryonic sex determination, gDNA isolated from E15.5 tail biopsies from 45 embryos was analyzed using both endpoint PCR and SYBR Green–based qPCR. Conventional PCR using *PRSSLY* and *KDM5C/D* primers yielded fully matching sex assignments across all samples (**Figure 3A**), demonstrating that *PRSSLY* amplification alone could serve as a reliable marker for male identification. Notably, a prominent 475 bp band was detected in several samples classified as female based on *KDM5C/D* (e.g. pups 5, 7, and 9, **Figure 3A**), alongside the characteristic faint nonspecific higher bands (∼1.5 kB) observed in female samples (**Figure 1C, 3A**).

**Figure 3.**
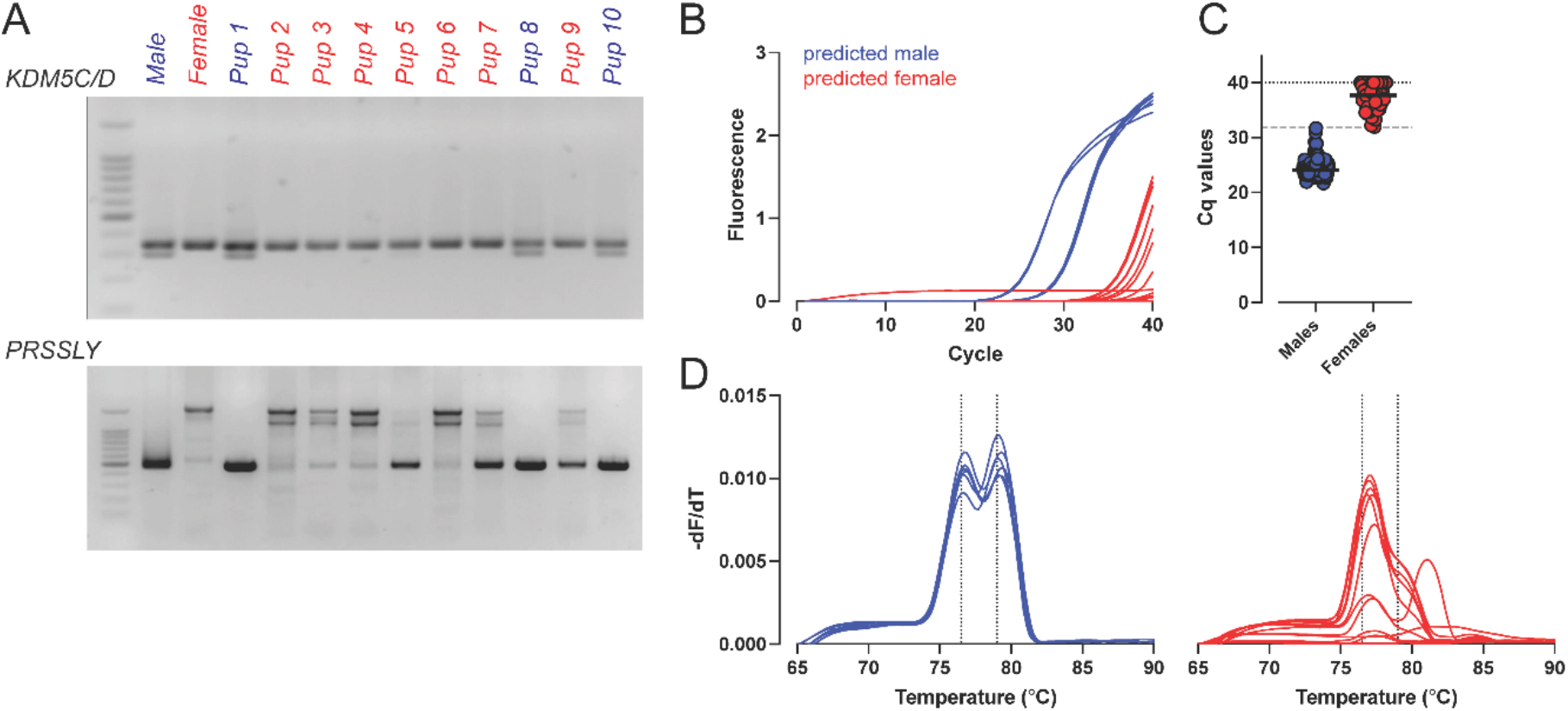
Validation of *PRSSLY-*based sex identification in embryonic tail biopsies at E15.5. (**A**) Representative endpoint PCR gels using tail-derived gDNA from E15.5 embryos, assayed with *KDM5C/D(*sex-control primers) and *PRSSLY*. Ten embryonic samples are shown, along with one adult male and one adult female as controls. (**B, D**) Real-time amplification curves (B) and corresponding melting curves (D) of *PRSSLY* in the same samples. Dotted vertical lines indicate the peak melting temperatures for the male-specific amplicon at 76.5 °C and 79.0 °C. (**C**) Cq values of all analysed embryonic samples (n = 209, 124 predicted males, 85 predicted females). The dashed horizontal line indicates the threshold Cq value used for gender assignment (Cq = 31.82), defined by ROC analysis (sensitivity = 1, specificity = 1, AUC = 1).

In contrast, qPCR provided clear and unambiguous discrimination between male and female samples. Male samples consistently showed early amplification and a distinct melt peak corresponding to the *PRSSLY*-specific amplicon at 76.5 °C and 79.0 °C, whereas all female samples had delayed amplification, displaying only background, nonspecific signals and no melt peak at 79.0 °C (**Figure 3B, D**).

Based on Cq values, ROC analysis identified a threshold of 31.82 (n = 209), which correctly segregated male and female samples with 100% sensitivity and 100% specificity (AUC = 1), with no overlap between groups (**Figure 3C**). These results demonstrate that *PRSSLY*-based qPCR offers a robust and highly discriminative method for embryonic sex identification, even in cases where conventional PCR produces ambiguous results.

We finally analyzed 417 placental samples collected from 39 dams using our SYBR Green– based *PRSSLY* melt-curve assay. Among these, 171 samples displayed a distinct melting peak and were classified as male, while 223 samples were negative, corresponding to female placentas (**Figure 4A-C**). An additional 23 samples exhibiting low amplitudes but detectable peaks were reanalyzed by agarose gel electrophoresis using targeted amplification in an independent PCR setup and confirmed as male. In parallel, 46 predicted female samples were also analyzed by gel electrophoresis, all yielding only faint, nonspecific products, validating their classification as female placentas. Notably, sex genotyping using the *KDM5C/D* primer set produced inconsistent melt peaks in placental gDNA, indicating its limited reliability for accurate sex determination in embryonic tissue (**Supplementary Figure 2**).

**Figure 4.**
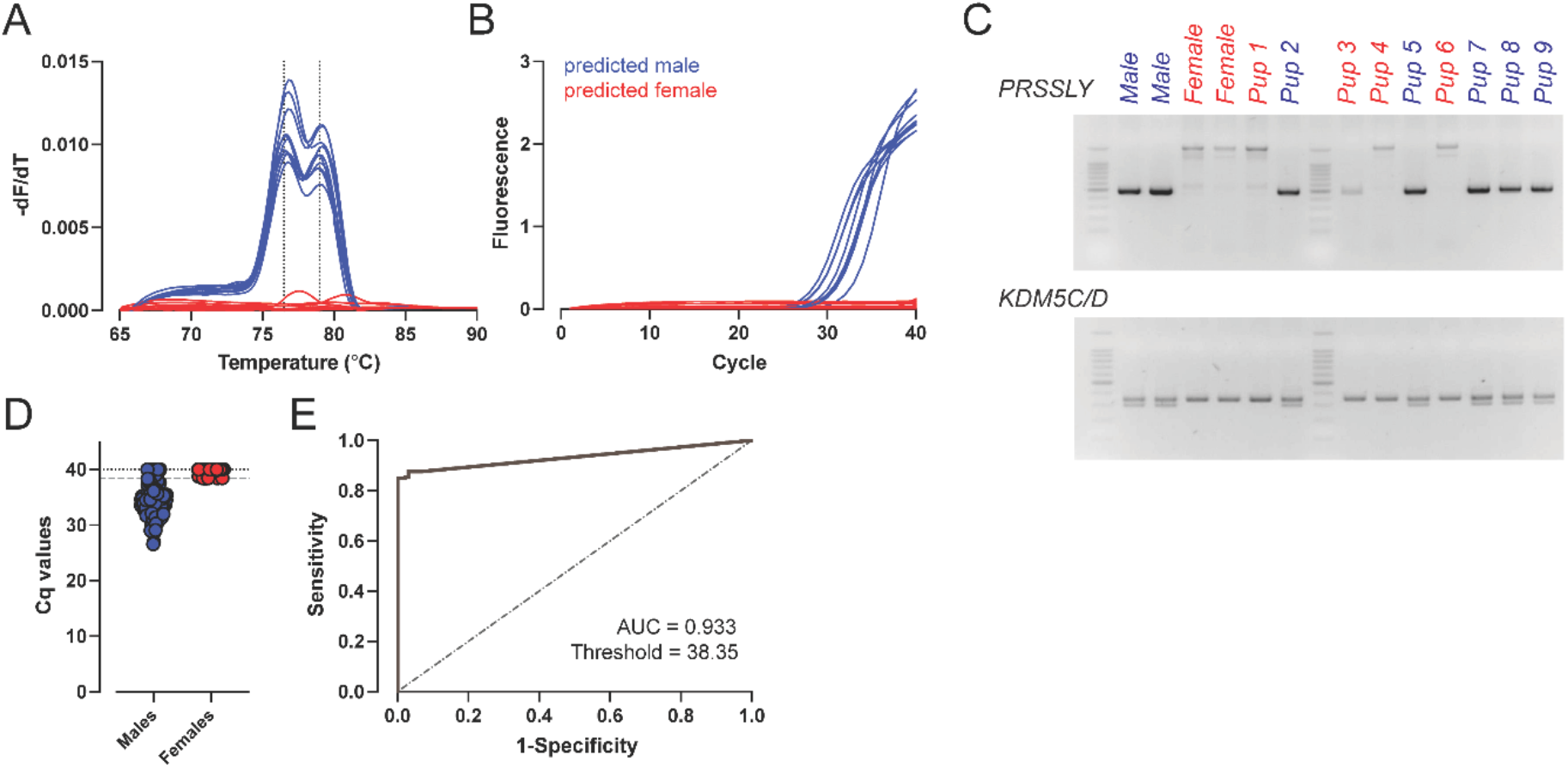
Sex determination by *PRSSLY*-based melt-curve assay using placental tissue. (**A, B**) Melting curves (A) and amplification curves (B) obtained from placenta-derived gDNA at E15.5 from one representative litter (n = 9 pups). Dotted vertical lines in A indicate peak melting temperatures of the male-specific *PRSSLY* amplicons at 76.5 °C and 79.0 °C. (**C**) Representative endpoint PCR gel electrophoresis using *PRSSLY* and *KDM5C/D* primer sets on samples corresponding to (A, B). Adult male and female gDNA were included as controls. (**D, E**) Distribution of Cq values for all analyzed placental samples (n = 417; 194 predicted males, 223 predicted females). The dashed horizontal line in (D) indicates the Cq cutoff (Cq = 38.35) that best discriminates male and female samples, as determined by ROC analysis (E) (AUC = 0.933, sensitivity = 0.85, specificity = 1.00).

Receiver operating characteristic (ROC) curve analysis using Cq values from all 417 samples identified an optimal discriminative threshold of Cq < 38.35 for male prediction, corresponding to a sensitivity of 85% and a false-positive rate of 0% as calculated by ROC analysis (**Figure 4D, E**).

Altogether, this work establishes a reliable molecular approach for sex determination in Syrian hamster placental and embryonic tissues, based on the detection of the Y-linked *PRSSLY* gene. The assay expands the available toolkit for genetic studies in this species, complementing recent advances in chromosome-scale genome assemblies derived from both female and male individuals [14-16]. These genomic resources eventually enable the systematic identification of male-specific loci through comparative and cross-species analyses, facilitating the development of robust diagnostic markers.

To note, despite a generally high >85% specificity, our *PRSSLY-*based assay occasionally produces low-amplitude peaks in male samples, which may complicate interpretation in placental tissues with low fetal DNA content. To further increase reliability, the method could be combined with amplification of an X-linked gene product, such as *ZFX*, or basically any autosomal gene providing an internal control for DNA integrity and qPCR performance. In addition, dubious samples could be retested by gel electrophoresis as demonstrated here (**Figure 3A, 4C**). Together, these refinements will support the accurate and reproducible genotyping of sex in developmental and reproductive studies using Syrian hamsters.

## Acknowledgments

The authors are grateful to Femke Vanzeebroeck, Shana Robberechts, Valentijn Vergote and the staff of the animal facility at the Rega Institute (KU Leuven) for their support in husbandry and time mating of hamsters.

## Funding

Flemish Research Foundation (FWO) grant G0H3120N (KD)

FWO Excellence of Science grant 40007527 (KD)

KU Leuven Global Seed Fund GSF/25/074 (YAA, KD)

European Union MSCA4Ukraine fellowship, grant number 101110724 (YK)

European Union Horizon Europe Programme, grant agreement number 101137459 (KD)

Views and opinions expressed are however those of the author(s) only and do not necessarily reflect those of the European Union, HADEA or the MSCA4Ukraine Consortium. Neither the European Union nor the granting authority, nor the MSCA4Ukraine Consortium as a whole nor any individual member institutions of the MSCA4Ukraine Consortium can be held responsible for them.

## Author Contributions

Conceptualization, Y.A.A., K.D.;

methodology, Y.K., V.N., Y.A.A., K.D.;

formal analysis, Y.K., Y.A.A.;

investigation, Y.K., L.P., V.N.;

resources, Y.A.A., K.D.;

data curation, Y.K., Y.A.A.;

writing—original draft preparation, Y.A.A., K.D.;

writing—review and editing, all authors;

visualization, Y.K., Y.A.A.;

supervision, Y.A.A., K.D.;

project administration, Y.A.A., K.D.;

funding acquisition, Y.K., Y.A.A., K.D.

All authors have read and agreed to the published version of the manuscript.

## Data and materials availability

All data supporting the findings in this study are available from the corresponding author upon reasonable request.

## SUPPLEMENTARY INFORMATION

### Sex determination of Syrian hamster pups using placental tissues

Yana Kumpanenko, Lindsey Piessens, Victor Neven, Kai Dallmeier, and Yeranddy A. Alpizar

## Supplementary Figures

**Supplementary Figure 1.**
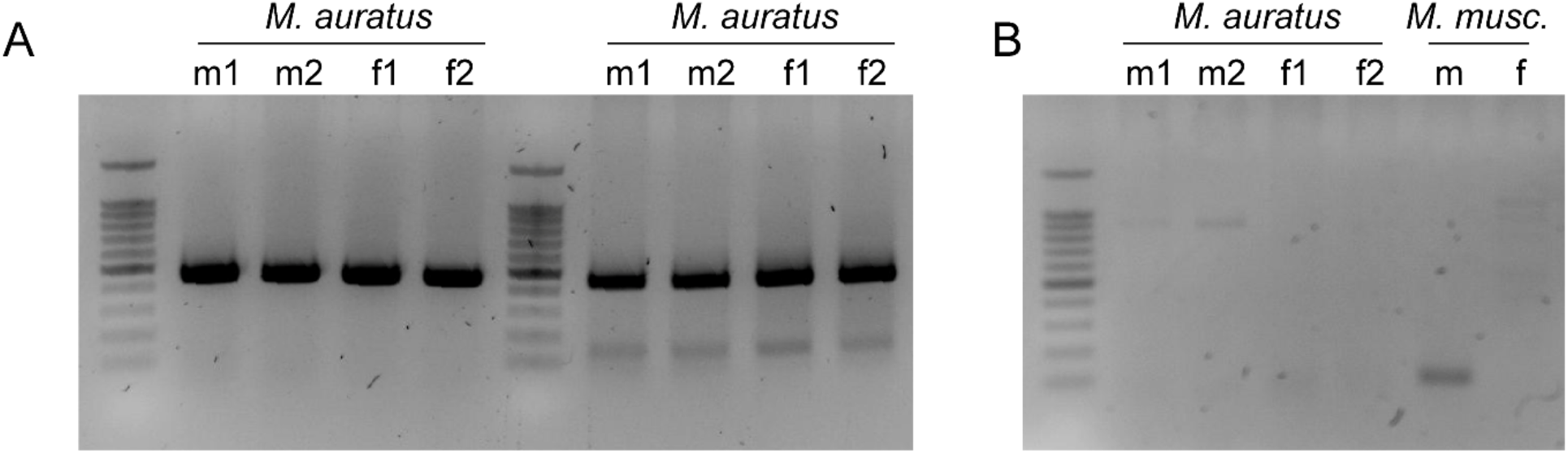
Lack of sex-specific amplification in Syrian hamsters using universal *SRY/TDY* primers. **(A)** PCR amplification using two primer sets targeting the *SRY* gene. The SRYA-5 primer set is shown on the left side of the gel, and the SRY_Mm primer set on the right. Both reactions were performed using genomic DNA isolated from ear biopsies of adult Syrian hamsters. **(B)** PCR amplification using the maTdy primer set. Reactions were performed using genomic DNA isolated from ear biopsies of adult Syrian hamsters and adult mice. Across both panels, the universal primer sets did not produce sex-specific PCR products in Syrian hamsters.

**Supplementary Figure 2.**
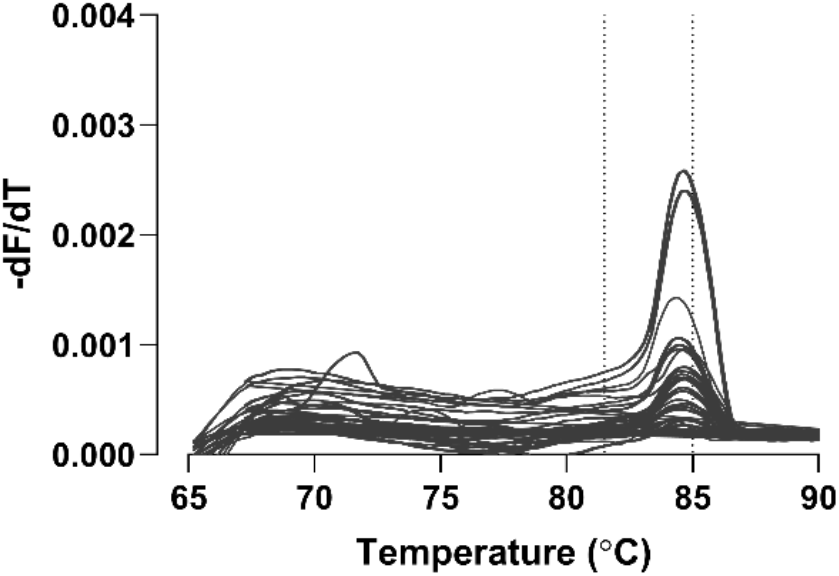
Melt-curve analysis of placental gDNA using the Kdm5c primer set. Melt curves generated from placental gDNA using Kdm5c primers. Samples correspond to the same litter shown in Figure 4 A-C. Dotted vertical lines indicate the expected male-specific melt peak at 81.5 °C and the common peak at 85.0 °C, as defined in [10].

